# Glutamine alters the response to stress in mice with diet-induced obesity in a sex-dependent manner

**DOI:** 10.1101/2025.11.21.689652

**Authors:** Candice Lefebvre, Adam Tiffay, Charles-Edward Breemeersch, Virginie Dreux, Christine Bôle-Feysot, Charlène Guérin, Jonathan Breton, Ludovic Langlois, Elise Maximin, Magali Monnoye, Pierre Déchelotte, Véronique Douard, Alexis Goichon, Moïse Coëffier

## Abstract

**Rationale:** Patients with class III obesity often suffer from irritable bowel syndrome (IBS) while obesity and IBS share common pathophysiological mechanisms such as altered intestinal barrier function and gut microbiota dysbiosis. Oral glutamine (Gln) supplementation previously showed beneficial effects on gut barrier function in a sex-dependent manner and reduced abdominal pain in IBS patients. Thus, we assessed the sex-dependent response to an oral Gln supplementation in mice with diet-induced obesity and subjected to a chronic restraint stress to mimic IBS.

**Methods:** Male (M) and female (F) C57BL/6 mice received a high fat diet (HFD; 60% kcal from fat) during 14 weeks (W14) and were subjected or not to a restraint stress (S) for the 4 last days. From W12, mice received or not Gln in drinking water (2g/kg/day; n=12/group). Plasma corticosterone, body composition, glucose tolerance (OGTT), intestinal permeability, inflammatory markers in colonic and white adipose tissues, cecal microbiota and short-chain fatty acid (SCFA) composition have been assessed. Within each sex, groups were compared by Kruskal-Wallis test or a 1-way ANOVA test.

**Results:** In M-HFD mice, chronic restraint stress was associated with a better glucose tolerance (–15,46% AUC) and a reduced fasting glycemia that was not observed in F-HFD mice. Gln partially prevented body weight loss, reduced plasma resistin, plasma corticosterone and colonic permeability in female stressed obese mice. In male stressed obese mice, Gln limited lean mass loss, reduced colonic permeability and Ccl2 mRNA level in the subcutaneous adipose tissue. Chronic restraint stress and Gln modified cecal microbiota in both sexes but cecal SCFA composition only in male mice. In particular, stress induced an increased abundance of *Pseudomonadota* in male mice that was partially restored after Gln supplementation. In addition, cecal total SCFA were reduced in Gln-supplemented stressed HFD male mice compared to unstressed HFD mice. In female HFD mice, stress associated to Gln supplementation increased the abundance of *Thermodesulfobacteriota* and reduced colonic expression of Cxcr3 mRNA.

**Conclusions:** Chronic restraint stress has beneficial effects on glycemia control in male HFD mice without additive effects of Gln supplementation. By contrast, Gln reduces stress-induced corticosterone level and body weight loss only in females. These data, as well as the differential impact of Gln on intestinal permeability and gut microbiota according to the sex, deserves further investigations to decipher the underlying mechanisms.

## Introduction

Obesity, defined by abnormal accumulation of fat and a body mass index (BMI) ≥30 kg/m², is a major public health issue (1,2) and is often associated with metabolic complications, *e.g.* type 2 diabetes, dyslipidemia, and increases the risk to develop chronic diseases. Obesity is a multifactorial pathology, linked to genetic, metabolic and environmental factors.

The role of gut microbiota and intestinal barrier function in obesity has gained significant attention, particularly regarding their regulation of low-grade systemic inflammation and glycemia control dysregulations (3). While studies in animal models of obesity have consistently described disruptions in intestinal barrier function (4–6), data in humans remain controversial (7). Beyond the association between obesity and intestinal barrier function, recent studies have also suggested a link between obesity and functional gastrointestinal disorders (FGIDs), now referred to as disorders of the gut-brain interaction (DGBI) (8,9). The prevalence of irritable bowel syndrome (IBS), a DGBI, was increased in European patients with grade III obesity (BMI > 40) to achieve 7.7% compared to 4.2% in the general population (10). Conversely, in another retrospective study, grade I and II obesity appeared to be associated with reduced risk of IBS (9). Interestingly, obesity and IBS share common pathophysiological mechanisms: gut dysbiosis, disruption of gut barrier function and low-grade inflammatory state. We previously observed that the expression of occludin, a tight junction protein, was more severely reduced in IBS patients with BMI >30 kg/m² compared to non-obese IBS patients (11). Similarly, gut barrier disruption was more pronounced in obese mice compared to lean mice in response to stress (12) a common trigger of both IBS and obesity (13).

Glutamine (Gln), the preferred substrate of rapidly dividing cells, is involved in several metabolic processes such as lipolysis, gluconeogenesis and protein synthesis (14). It appears as a good candidate to limit obesity-associated intestinal disorders. Indeed, even if we recently reported that Gln supplementation failed to improve colonic paracellular permeability in obese mice (15), pilot studies showed that Gln supplementation in subjects with obesity was able to reduce fat mass, waist circumference, circulating LPS (16–18), and to restore *Bacillota/Bacteroideta* ratio (19). Gln supplementation also improved glycemia homeostasis in reducing insulin sensitivity (18) and glucose tolerance (15) in male rodent fed with a high fat diet (HFD), but not in females. Stress, which is known to play a significant role in the development and exacerbation of IBS symptoms, has been shown to activate the hypothalamic–pituitary–adrenal (HPA) axis and to disturb the gut barrier function (20). Interestingly, Gln supplementation prevented (i) colonic hyperpermeability in IBS-like mice submitted to water-avoidance stress (21) or chronic restraint stress (22), and (ii) reduced significantly stool frequency and abdominal pain in IBS patients with predominant diarrhea (23). Despite of these studies, the impact of Gln on IBS symptoms induced by stress in obese mice remains unknown.

Taking into account all these data, we aimed in the present study to assess the impact of oral Gln supplementation on the stress response induced in mice with diet-induced obesity by evaluating gut barrier function, inflammatory responses, gut microbiota and metabolic complications. Finally, as these alterations could be differently affected between males and females as highlighted by recent experimental studies (15,24–26), we thus performed the experiments both in male and female mice.

## Material and methods

### Animal experimentation

Six-week-old male (M) and female (F) C57BL/6J mice (Janvier Labs, Le Genest-Saint-Isle, France, n=48/sex) were acclimatized in a controlled environment (23°C with a 12 hours light-dark cycle) for 4 days with free access to water and a Standard Diet (SD, 3.34 kcal/g, 1314, Altromin, Germany). M– and F-mice were then fed with High Fat Diet for 14 weeks (HFD, 5.21 kcal/g; D12492i, Research Diet Inc., US). HFD supplied 60% kcal as fat, 20% kcal as carbohydrates and 20% kcal as proteins. At week 12 (W12), M– and F-mice were randomized into three groups (n=12/group): HFD (HFD-NS), HFD+stress (HFD-S) or HFD+Stress+Gln (HFD-S+Gln). From W12, mice received or not glutamine (Gln, G5792, Merck, Germany) during 15 days in drinking water to provide 2 g.kg^−1^ of body weight per day^−1^. The last 4 days of the experiments, mice were subjected to a repeated restraint stress. Body weight was measured weekly to follow the impact of HFD and Gln supplementation, as well as before and after stress procedures. Similarly, body composition was measured by EchoMRI (EchoMRI, Houston, US). The protocol was approved by the regional ethics committee (authorization on APAFIS #29283-2021012114574889 v5) and performed in accordance with the current French and European regulations.

### Chronic restraint stress

We employed the chronic restraint stress model (CRS) to induce IBS-like symptoms in mice: altered gut barrier function, gut dysbiosis, increased fecal number, and anxiety– and depression-like behaviors. We previously validated this approach in comparison to an acute restraint stress protocol (22). Mice were briefly anesthetized with isoflurane and placed in restraint cages (Bioseb®, Vitrolles, France) for two hours before returning to their home cage. The restraint sessions were repeated for four consecutive days (at the same hour) before sacrificing the animals. Control mice were kept in their home cage during the procedure.

### Oral glucose tolerance test (OGTT)

Oral glucose tolerance test (OGTT) was realized before (W12) and after Gln supplementation and stress procedures (W14) as previously described (26). Briefly, after an overnight fasting followed by an oral gavage of glucose (1 g. kg^-1^ of body weight; G8769, Merck), glycemia was measured at 0, 15, 30, 45, 60 and 90 minutes using a glucometer (Accu Check Guide, Roche, Switzerland).

### Intestinal permeability

Whole intestinal permeability was first measured *in vivo* at the end of the protocol (week 14 – W14). Briefly, After a 2 hours fasting period and 3 hours before euthanasia, mice received a FITC-dextran solution (4 kDa, 40 mg/mL, Merck) by oral gavage at 10 µL/g of body weight (26). After Gln supplementation and stress procedures (W14), colonic paracellular permeability was also evaluated *ex vivo* by measuring fluxes of Alexa Fluor 680-dextran (3 kDa, 0.02 mg/mL, Merck) and Lucifer Yellow (457 Da, 2.5 mg/mL, Merck) across colonic mucosa mounting in Ussing Chambers (Harvard Apparatus, US) during 3 hours at 37°C. Lucifer yellow is generally used as a probe for investigating the three major paracellular pathways (pore, leak, and unrestricted) whereas Alexa Fluor 680-dextran is a leak and unrestricted pathway probe. The fluorescence intensities of FITC-dextran (excitation at 485 nm, emission at 535 nm) in plasma or Alexa Fluor 680-Dextran (excitation at 665 nm, emission at 710 nm) and Lucifer Yellow (excitation at 428 nm, emission at 540 nm) in serosal medium were measured with the Spark multimode microplate reader (Tecan, UK).

### Tissue sampling

At W14, mice were anesthetized by i.p. injection of ketamine/xylazine solution (100 and 10 mg/kg, respectively). Blood samples were collected, centrifuged for 15 min at 3000*xg* at 4°C and plasma was frozen at –80°C. Subcutaneous and perigonadal adipose tissue depots and colon were collected, washed with ice-cold PBS, immediately frozen in liquid nitrogen and stored at –80°C until analysis. Cecal content was collected to evaluate microbiota and short-chain fatty acids.

### Plasma assays

Plasma levels of insulin, leptin, adiponectin, resistin, GLP-1 and pro-inflammatory cytokines (IL-6 and TNFα) were assessed using Bio-Plex Pro assays (BioRad Laboratories, Marnes-la-Coquette, France) and Luminex technology as previously described (26). Plasma urea, total proteins, alanine aminotransferase (ALAT), aspartate aminotransferase (ASAT), gamma-glutamyltransferase (GGT), total cholesterol and triglycerides (TG) concentrations were determined using a Catalyst One device (IDEXX Laboratories, Westbrook, ME, US).

### Colonic RNA extraction and RT-qPCR

Extraction of total colonic mRNAs, quantification and reverse transcription were performed as previously described (26). qPCR were realized by using the SYBR Green technology on a QuantStudio 12K Flex real-time PCR system (Life Technologies, Carlsbad, CA, US) targeting 44 markers of interest mainly involved in inflammatory response and gut barrier function. The mean of ACTB, β2M and GAPDH genes were used as reference. Gene-specific primers were previously detailed (26).

### Short chain fatty acid analysis

Twenty to fifty grams of frozen cecal contents were used to analyzed the short-chain fatty acids (SCFAs) composition using gas chromatography (Agilent 7890B gas chromatograph, Agilent Technologies, Les Ulis, France) as previously described (26).

### Cecal microbiota analysis

Total DNA was extracted from cecal content samples with PowerFecal Pro DNA Kit (Qiagen, France) as described previously (26) and the V3–V4 hypervariable region of the bacterial 16S rDNA was amplified by PCR using Phanta Max Super-Fidelity DNA Polymerase (Vazyme, PRC) (95°C for 3 min and then 35 cycles at 95°C for 15 sec, 65°C for 15 sec, and 72°C for 1 min before a final step at 72°C for 5 min) and the following primers: PCR1F_460: CTTTCCCTACACGACGCTCTTCCGATCTACGGRAGGCAGCAG and PCR1R_460: GGAGTTCAGACGTGTGCTCTTCCGATCTTACCAGGGTATCTAATCCT. The purified amplicons were sequenced using Miseq sequencing technology (Illumina) at the GeT-PLaGe platform (Toulouse, France). The resulting sequences were analysed using R (Team, 2019) workflow combining dada2 v.1.2830 and FROGS 4.0.131 software. Forwards and reverse reads were processed as described previously to obtained the amplicon sequence variants (ASV) (26). The resulting sequences were saved in a sequence table and the ASVs were then processed according to FROGS v4.0.1 guidelines and were then assigned to species using FROGS affiliation with 16S REFseq Bacteria.

Subsequent analysis were done using the phyloseq R packages (27). Samples were rarefied to even sampling depths before computing within-samples compositional diversities (observed richness, Chao1, Shannon and InvSimpson index) and between-samples compositional diversity (Bray-Curtis; Jaccard and Unifrac). Raw, unrarefied ASV counts were used to produce relative abundance graphs and to find ASV with significantly different abundances HFD-S and HFD-S-Gln in both male and female mouse groups.

### Adipose tissue RNA extraction and RT-qPCR

Extraction of perigonadal and subcutaneous white adipose tissue (WAT) mRNAs, quantification and reverse transcription were performed as previously described (26). qPCR were realized by using the SYBR Green technology on a Bio-Rad CFX96 real-time PCR system (Bio-Rad Laboratories). The mean of β2M and RPS18 genes was used as reference. Gene-specific primers were previously detailed (26).

### Statistical Analysis

Data were analyzed with GraphPad Prism 8.3 (GraphPad Software Inc., San Diego, CA, USA) and expressed as the mean ± standard error of the mean (SEM). Within each sex, groups were compared by a nonparametric Kruskal-Wallis test followed by Dunnett’s multiple comparison tests. To compare glycemia changes during the OGTT, two-way ANOVA tests were performed (Time x Groups) followed by Bonferroni post-tests. Alpha diversity index (Observed species, Chao1, Shannon and InvSimpson) were analyzed using Kruskal-Wallis test. A permutational multivariate analysis of variance (PERMANOVA) test was performed on the Bray–Curtis, Jaccard and Unifrac matrices using 9999 random permutations and at a significance level of 0.05. Phylum and family relative abundances were compared using Kruskal-Wallis test followed by Dunn’s post hoc test. For all, a value of p<0.05 was considered significant. A negative binomial model was fit to each ASV, using DESeq2 (28) with default parameters, to estimate abundance log2-fold changes. P-values were corrected for multiple testing using the BH procedure to control the False Discovery Rate (FDR) and significant ASVs were selected based on effect size [fold change (FC) > 4 or FC < 1/4], adjusted p-value (<0.05) and prevalence (more than 30 copies per sample in at least 25% of the samples).

## Results

### Impact of the stress and Gln on body weight and metabolic parameters

In both M-HFD and F-HFD mice, chronic restraint stress induced an increase in plasma corticosterone levels (Table 1) and a loss of body weight (Fig. 1A) compared to unstressed mice, validating this procedure as a method to induce stress in obese mice. Interestingly, Gln supplementation partially prevented the increase in plasma corticosterone levels (Table 1) and body weight loss (Fig. 1A) in F-HFD-S mice but had no impact in M-HFD-S mice even if fat mass tended to decrease after Gn supplementation in M-HFD-S mice (Fig. 1B, p=0.0521) but not in F-HFD mice. In addition, only M-HFD mice showed a significant reduction in lean mass following stress, which was not significant in Gln-supplemented group (Fig. 1B).

**Fig. 1:**
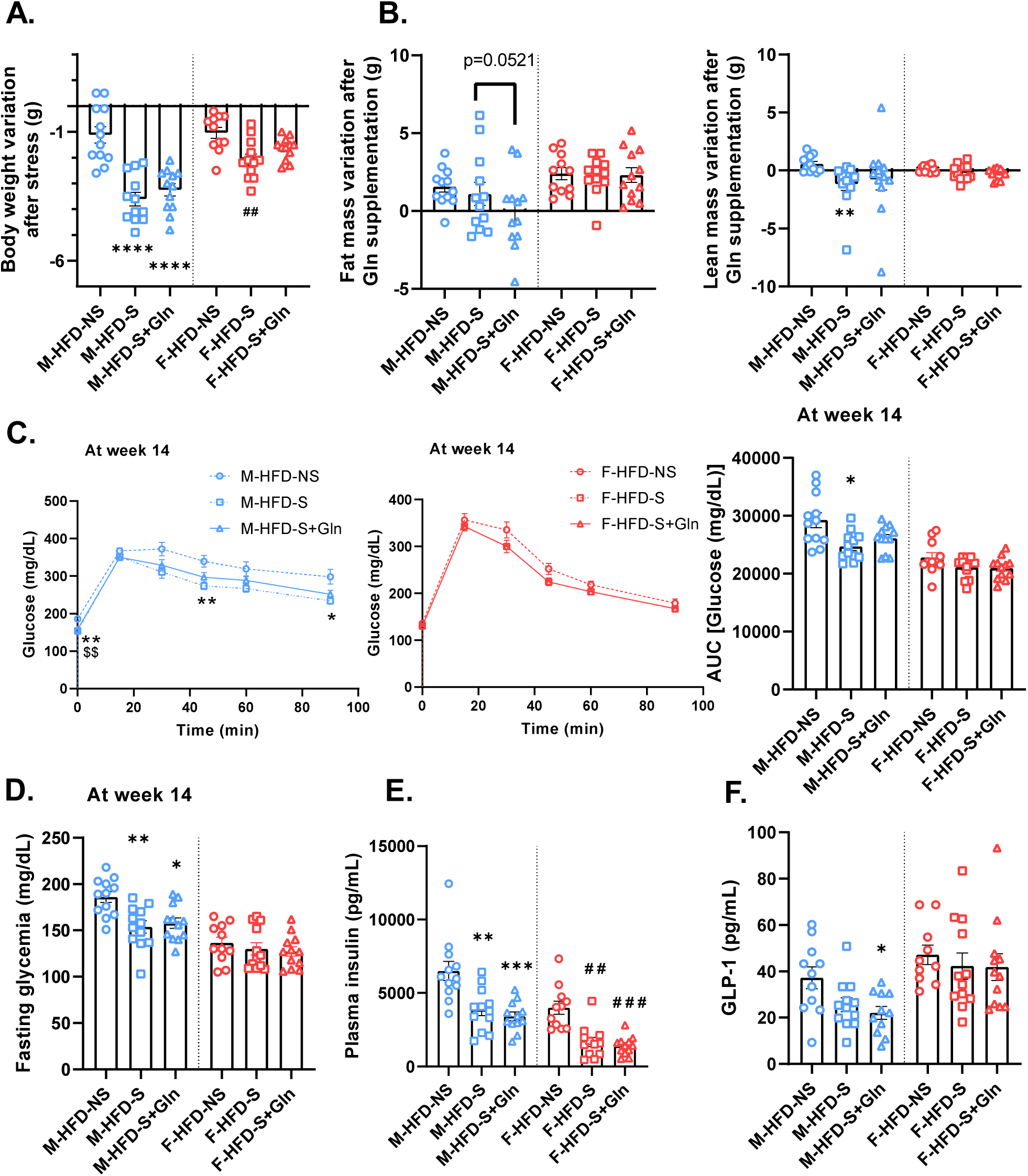
Effects of glutamine (Gln) supplementation on plasma corticosterone level, body weight, lean mass and glycemia control in obese male and female mice subjected to stress. Body weight changes induced by stress (A), fat and lean mass changes induced Gln supplementation (B) and oral glucose tolerance test (OGTT, C), fasting glycemia (D), insulin (E) and GLP-1 (F) plasma levels assessed at week 14 (W14) in male (M) and female (F) mice fed with high fat diet (HFD) and receiving or not glutamine (Gln) in drinking water from W12 to W14. The last 4 days of the experiments, mice were subjected to a repeated restraint stress (S). For A, B, C (AUC), D, E et F, results were compared by Kruskal-Wallis test followed by Dunnett’s multiple comparison tests. For C (glucose concentrations), results were compared by two-way ANOVA (time x groups) test followed by Bonferroni post-test. Data are expressed as mean ± sem (n=12/group). \**p*<0.05, \*\**p*<0.01, \*\*\**p*<0.001, \*\*\*\**p*<0.0001 M-HFD-S or M-HFD-S+Gn *vs* M-HFD-NS; #*p*<0.05, ##*p*<0.01, ###*p*<0.001 F-HFD-S or F-HFD-S+Gn *vs* F-HFD-NS; $$ *p*<0.01 M-HFD-S+Gn *vs* M-HFD-NS. AUC, Area under the curve.

**Table 1:** Plasma levels of metabolic markers in male and female obese mice subjected to stress and supplemented or not with glutamine.

| Markers | Experimental groups |  |  |  |  |  |
| --- | --- | --- | --- | --- | --- | --- |
|  | Male mice (M) |  |  | Female mice (F) |  |  |
|  | HFD-NS | HFD-S | HFD-S+Gln | HFD-NS | HFD-S | HFD-S+Gln |
| ALAT (U/L) | 88.75±10.56 | 137.8±19.69 | 130.80±16.79 | 60.00±2.96 | 75.08±5.19 | 91.50±6.07<sup>###</sup> |
| ASAT (U/L) | 218.3±13.62 | 273.0±20.18 | 238.8±18.70 | 200.2±11.93 | 228.8±18.05 | 239.0±23.50 |
| Cholesterol (g/L) | 1.22±0.053 | 1.24±0.068 | 1.19±0.087 | 0.67±0.034 | 0.61±0.051 | 0.59±0.049 |
| Corticosterone (ng/mL) | 12.21±1.87 | 68.57±11.62<sup>****</sup> | 59.08±8.15<sup>***</sup> | 51.35±6.63 | 150.80±29.19<sup>##</sup> | 94.66±12.18 |
| GGT (U/L) | 6.25±0.93 | 4.42±0.81 | 5.67±0.83 | 7.91±1.22 | 5.92±0.88 | 8.67±2.59 |
| Total protein (U/L) | 49.58±1.22 | 55.00±1.14<sup>**</sup> | 51.67±0.94 | 46.73±0.84 | 50.25±0.93 | 52.83±1.18<sup>##</sup> |
| Triglycerides (g/L) | 0.89±0.043 | 0.85±0.040 | 0.83±0.029 | 0.73±0.029 | 0.75±0.040 | 0.83±0.034 |
| Urea (g/L) | 0.31±0.017 | 0.27±0.024 | 0.29±0.073<sup>*</sup> | 0.38±0.019 | 0.33±0.026 | 0.31±0.011 |

Results were compared by Kruskal-Wallis test followed by Dunnett's multiple comparison tests. Data are expressed as mean ± sem (n=12/group). \* $p<0.05$ , \*\* $p<0.01$ , \*\*\* $p<0.001$ , \*\*\*\* $p<0.0001$ M-HFD-S or M-HFD-S+Gn vs M-HFD-NS; # $p<0.05$ , ## $p<0.01$ F-HFD-S or F-HFD-S+Gn vs F-HFD-NS. ALAT: alanine aminotransferase, ASAT: aspartate aminotransferase, GGT: gamma-glutamyltransferase.

In male mice, stress altered the regulation of glycemia homeostasis. Indeed, M-HFD-S mice exhibited improved tolerance during the OGTT compared to M-HFD-NS mice (Fig. 1C), which was associated with reduced fasting glycemia and insulinaemia (Fig. 1D and E). Although, Gln supplementation did not affect fasting glycemia and insulinaemia, it blunted the reduction in glucose AUC during OGTT. Additionally, a trend toward decreased plasma GLP1 levels was observed in stressed male mice fed with HFD, which reached significance in the Gln-supplemented group only (Fig. 1F). In F-HFD mice, stress did not affect glucose tolerance (Fig. 1C) or fasting glycemia (Fig. 1D), but it did reduce fasting insulinaemia (Fig. 1E). Gln supplementation had no effect on these glycemia parameters in female mice.

We then measured inflammatory and metabolic parameters in plasma samples. In both male and female mice, plasma levels of proinflammatory cytokines IL-6 and TNFα were not significantly modified (data not shown). As shown in Table 1, neither stress nor Gln supplementation substantially modified dyslipidemia (cholesterol, triglycerides), protein metabolism (urea, total protein), or hepatic markers (ALAT, ASAT, γGT). Gln supplementation associated to stress increased plasma total protein and ALAT levels in F-HFD mice and reduced plasma urea levels in M-HFD mice.

### Impact of the stress and Gln on the adipocyte response

In perigonadal and subcutaneous adipose tissues, we were not able to show difference in the mRNA levels for leptin, adiponectin and resistin in both male and female mice (Supplemental Fig. S1). We then measured adipokine concentrations in the plasma. In male mice, plasma leptin and adiponectin concentrations remained unchanged, while resistin concentrations were lower in the stressed groups (Fig. 2). In female mice, stress did not affect significantly plasma leptin, adiponectin and resistin levels (Fig. 2), even if a trend for a decrease in resistin was observed (p=0.06). Gln supplementation did not affect plasma leptin while it is associated to an increase in adiponectin and a decrease in resistin levels in plasma samples in F-HFD mice (Fig. 2).

**Fig. 2:**
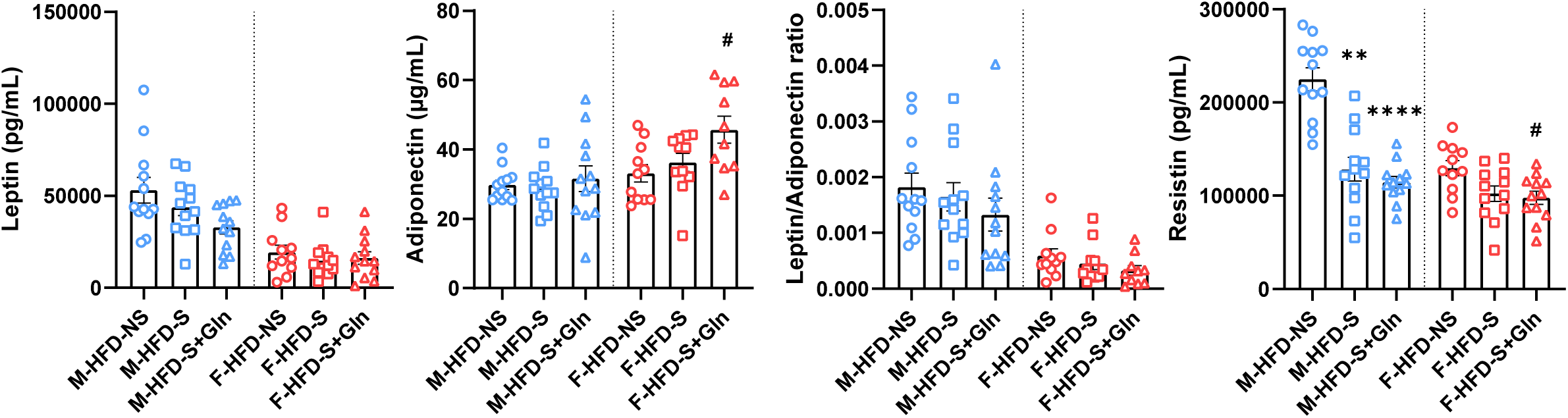
Effects of glutamine (Gln) supplementation on plasma adipokine levels in obese male and female mice subjected to stress. Plasma leptin, adiponectin and resistin concentrations were assessed at week 14 (W14) in male (M) and female (F) mice fed with high fat diet (HFD) and receiving or not glutamine (Gln) in drinking water from W12 to W14. The last 4 days of the experiments, mice were subjected to a repeated restraint stress (S). Results were compared by Kruskal-Wallis test followed by Dunnett’s multiple comparison tests. Data are expressed as mean ± sem (n=12/group). \**p*<0.05, \*\**p*<0.01, \*\*\**p*<0.001, \*\*\*\**p*<0.0001 M-HFD-S or M-HFD-S+Gn *vs* M-HFD-NS; #*p*<0.05 F-HFD-S or F-HFD-S+Gn *vs* F-HFD-NS.

We also evaluated the mRNA levels for pro-inflammatory cytokines [IL-6, TNFα and Ccl2 (also called MCP-1)] in the perigonadal and subcutaneous fat tissues (Supplemental Fig. S1). Stress significantly reduced the mRNA levels for IL-6 in perigonadal adipocytes in male mice and the mRNA levels for Ccl2 in the subcutaneous adipose tissue in female mice. Gln supplementation was able to reduce Ccl2 mRNA levels in the subcutaneous adipose tissue in stressed male mice but had no effects in female mice.

### Impact of the stress and Gln on intestinal response

First, we evaluated the whole intestinal permeability using oral gavage with FITC-dextran (4kDa). Stress did not increase whole intestinal permeability in either male and female mice compared to HFD-NS mice (Fig. 3A). However, Gln supplementation increased whole intestinal permeability in stressed male mice compared to HFD-NS mice, an effect that was not observed in females (Fig 3A). Next, we examined colonic permeability using Lucifer Yellow (457 Da) and Alexa Fluor 680-D (3kDa) flux in Ussing chambers. Consistent with the whole intestinal permeability data, stress did not affect colonic permeability (Fig. 3A). Interestingly, Gln supplementation associated to stress significantly reduced colonic paracellular permeability both in M-HFD and F-HFD mice, except for Alexa Fluor 680-D in males. To clarify the data on the intestinal permeability, we assessed the mRNA expression levels of the tight junction proteins in the colonic mucosa. In accordance with Ussing chamber data, stress did not change tight junction protein mRNA expression in both sexes (Fig. 3B), excepted for claudin-5 (Cldn-5) mRNA levels. In stress situation, Gln enhanced mRNA levels of claudin-1 (Cldn-1) in male mice but not in female mice. By contrast, in stressed female mice, Gln reduced the mRNA expression of Cldn-12 and Ocln that was not observed in males. These data suggest that stress alone did not impact intestinal permeability in obese mice but that Gln supplementation associated to stress decreased colonic paracellular permeability in obesity context.

**Fig. 3:**
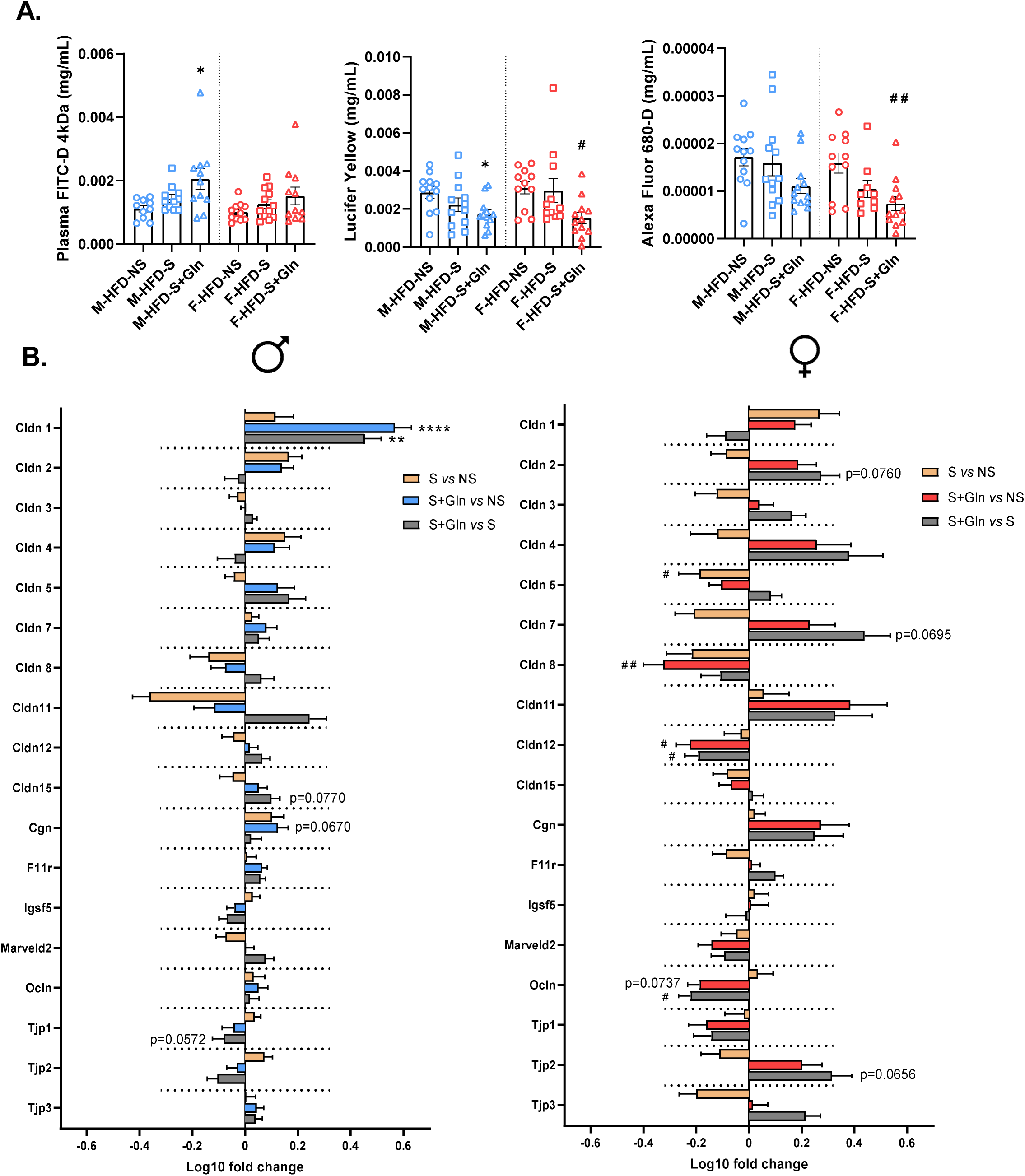
Effect of glutamine (Gln) supplementation on intestinal permeability and colonic gut barrier markers in obese male and female mice subjected to stress. Global intestinal permeability (FITC-D, **A**), colonic permeability (Lucifer Yellow and Alexa Fluor 680-D; **A**) and mRNA colonic levels of gut barrier markers (**B**) were assessed at week 14 (W14) in male (M) and female (F) mice fed with high fat diet (HFD) and receiving or not glutamine (Gln) in drinking water from W12 to W14. The last 4 days of the experiments, mice were subjected to a repeated restraint stress (S). Results were compared by Kruskal-Wallis test followed by Dunnett’s multiple comparison tests. For **B**, data are presented as ratio log10 [(HFD-S or HFD-S+Gln) / mean HFD-NS] and expressed as mean ± sem (n=12/group). \**p*<0.05, \*\**p*<0.01, \*\*\**p*<0.001, \*\*\*\**p*<0.0001 M-HFD-S or M-HFD-S+Gn *vs* M-HFD-NS; #*p*<0.05, ##*p*<0.01 F-HFD-S or F-HFD-S+Gn *vs* F-HFD-NS. *Cgn*: Cingulin, *Cldn1*: Claudin 1, *Cldn2*: Claudin 2, *Cldn3*: Claudin 3, *Cldn4*: Claudin 4, *Cldn5*: Claudin 5, *Cldn7*: Claudin 7, *Cldn8*: Claudin 8, *Cldn11*: Claudin 11, *Cldn12*: Claudin 12, *Cldn15*: Claudin 15, *F11r*: F11 receptor or JAM-A, *Igsf5*: Immunoglobulin superfamily member 5 or JAM-4, *Marveld2*: Marvel domain-containing protein 2 or Tricellulin, *Ocln*: Occludin, *Tjp1*: Tight junction protein 1 or ZO-1, *Tjp2*: Tight junction protein 2 or ZO-2; *Tjp3*: Tight junction protein 3 or ZO-3.

The TLR-mediated signaling pathways contributes significantly to obesity-associated inflammation by promoting production of proinflammatory cytokines. Consequently, we studied the TLR-2/4 pathways and some cytokines in colonic mucosa of HFD mice. Stress only induced a reduction of Cxcr3 mRNA levels in males compared to unstressed mice (Fig.4A). In female mice, stress did not affect cytokine mRNA levels but reduced Myd88 and Nod2 factors (Fig. 4B). Gln supplementation reduced the mRNA expression of Cxcr3 in F-HFD-S female mice, while it was the opposite for male mice.

**Fig. 4:**
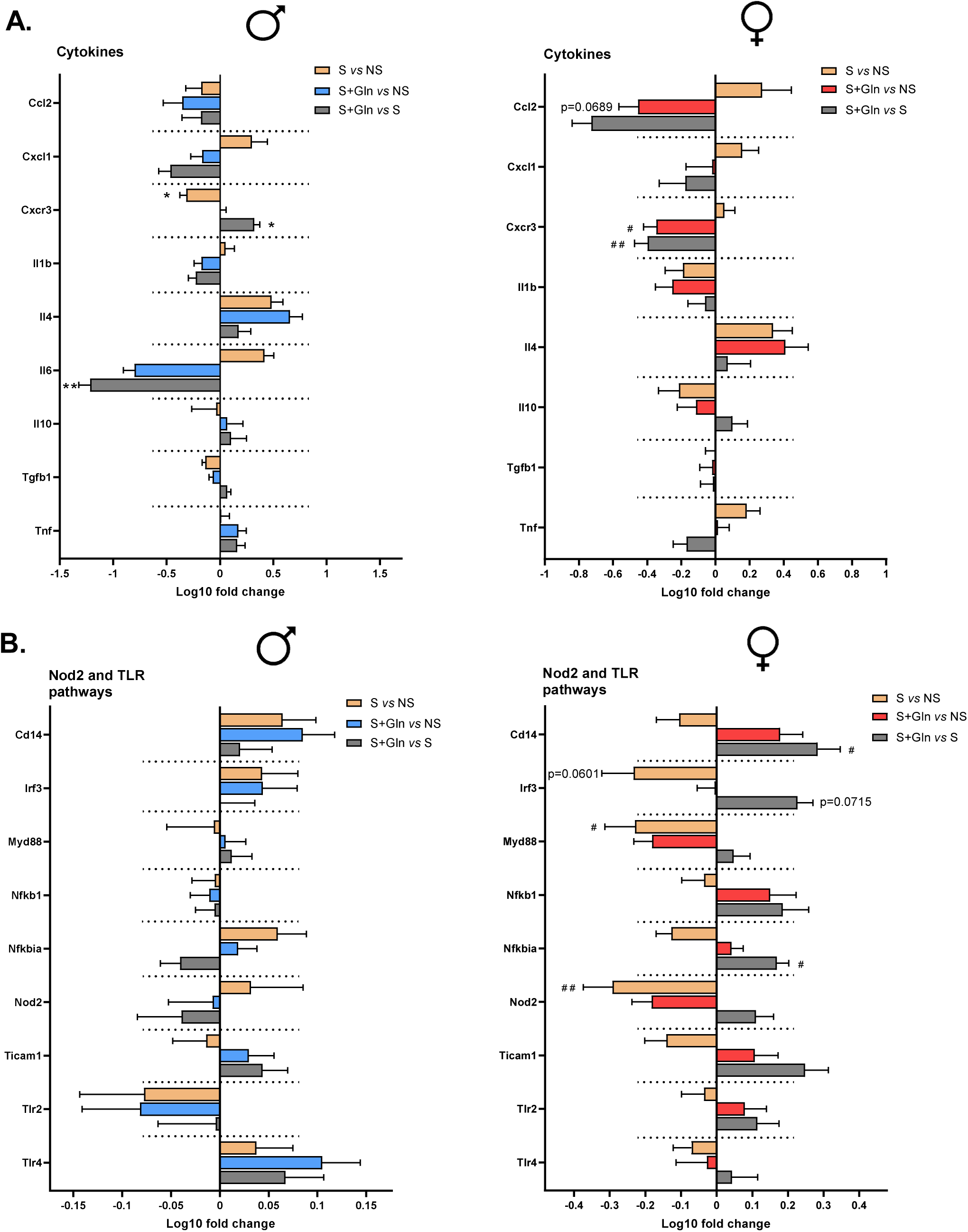
Effect of glutamine (Gln) supplementation on colonic gut barrier inflammatory markers in obese male and female mice subjected to stress. mRNA colonic levels of cytokines (**A**) and Nod2-TLR pathways markers (**B**) were assessed at week 14 (W14) in male (M) and female (F) mice fed with high fat diet (HFD) and receiving or not glutamine (Gln) in drinking water from W12 to W14. The last 4 days of the experiments, mice were subjected to a repeated restraint stress (S). Results were compared by Kruskal-Wallis test followed by Dunnett’s multiple comparison tests. Data are presented as ratio log10 [(HFD-S or HFD-S+Gln) / mean HFD-NS] and expressed as mean ± sem (n=12/group). \**p*<0.05, \*\**p*<0.01 M-HFD-S or M-HFD-S+Gn *vs* M-HFD-NS; #*p*<0.05, ##*p*<0.01 F-HFD-S or F-HFD-S+Gn *vs* F-HFD-NS. *Ccl2*: C-C motif Chemokine ligand 2 (or MCP-1), *Cd14*: CD14 antigen, *Cxcl1*: C-X-C motif chemokine ligand 1, *Cxcr3*: C-X-C motif chemokine receptor 3, *Il1b*: Interleukin 1 beta, *Il10*: Interleukin 10, *Il4*: Interleukin 4, *Il6*: Interleukin 6, *Irf3*: Interferon regulatory factor 3, *Myd88*: Myeloid differentiation primary response 88, *Nfkb1*: Nuclear factor kappa B1, *Nfkbia*: Nuclear factor kappa B inhibitor alpha, *Nod2*: Nucleotide-binding oligomerization domain-containing protein 2, *Tgfb1*: Transforming growth factor beta 1, *Ticam1*: TIR domain-containing adapter molecule 1, *TLR2*: Toll-like receptor 2, *TLR4*: Toll-like receptor 4, *Tnf*: Tumor necrosis factor alpha.

### Impact of the stress and Gln on cecal microbiota

Given the role of the intestinal microbiota in the adaptation of the intestinal epithelium to nutrients, we then analyzed the composition of the microbiota of the cecal contents (Fig. 5 and supplementary Fig. S3) and measured its concentrations of short-chain fatty acids (Fig. 6). We first examined gut microbiome diversity between the three groups in male and female mice. We found no significant differences between groups in either sex (Supplemental Fig. S2A). To assess differences in overall community structure, beta diversity was analyzed between the groups. Significant differences in community structure were observed in response to stress in both M-HFD and F-HFD mice, as measured by three beta diversity metrics: Jaccard, Bray-Curtis, and UniFrac (Supplemental Fig. S2B). However, these data suggest that Gln supplementation does not result in major changes in community structure. To broadly visualize the gut microbiome composition across groups, we examined the distribution of bacterial phyla and families. In male mice, at the phylum levels, stress induced an increase of *Pseudomonadota* that was restored after Gln supplementation (Fig. 5A). The abundance of other phyla was not changed. By contrast, in female mice, stress reduced the abundance of *Bacteroidota* but did not affect *Pseudomonadota* and other phyla. Gln supplementation did not affect *Bacteroidota* abundance in F-HFD-S mice (Fig. 5A). At the family level, under HFD and in agreement with previous studies (26), the most abundant families in both male and female mice (Fig. 5B) are *Lachnospiraceae*, *Desulfovibrionaceae*, *Oscillospiraceae*, and *Rikenellaceae*, which together represent more than 80% of all families. In male mice (Fig. 5C), stress reduced the abundance of *Lactobacillaceae*, while increasing the abundances of *Eggerthellaceae, Tannerellaceae, Sutterellaceae*, and *Clostridiaceae*. Notably, Gln supplementation restored *Sutterellaceae* abundance and increased the abundance of *Rikenellaceae* in stress situation. In female mice, stress was associated with an increase in *Eubacteriales_Family_XII_Incertae_Sedis*, which was fully restored by Gln supplementation (Fig. 5C). When combined with Gln supplementation, stress significantly reduced the abundances of *Rikenellaceae* and *Streptococcaceae*, while increasing those of *Lachnospiraceae* and *Desulfovibrionaceae*. Additionally, under stress conditions, Gln supplementation significantly increased the relative abundance of *Odoribacteraceae*. The Supplemental Fig. S3 shows the ASVs with a Log2 fold change > 2 or < –2 between HFD-S+Gln and HFD-S mice. In male mice, we identified 12 ASVs exhibiting a greater abundance in response Gln supplementation mainly belonging to *Lachnospiraceae*, *Peptostreptococcaceae* and *Clostridiaceae* families.

**Fig. 5:**
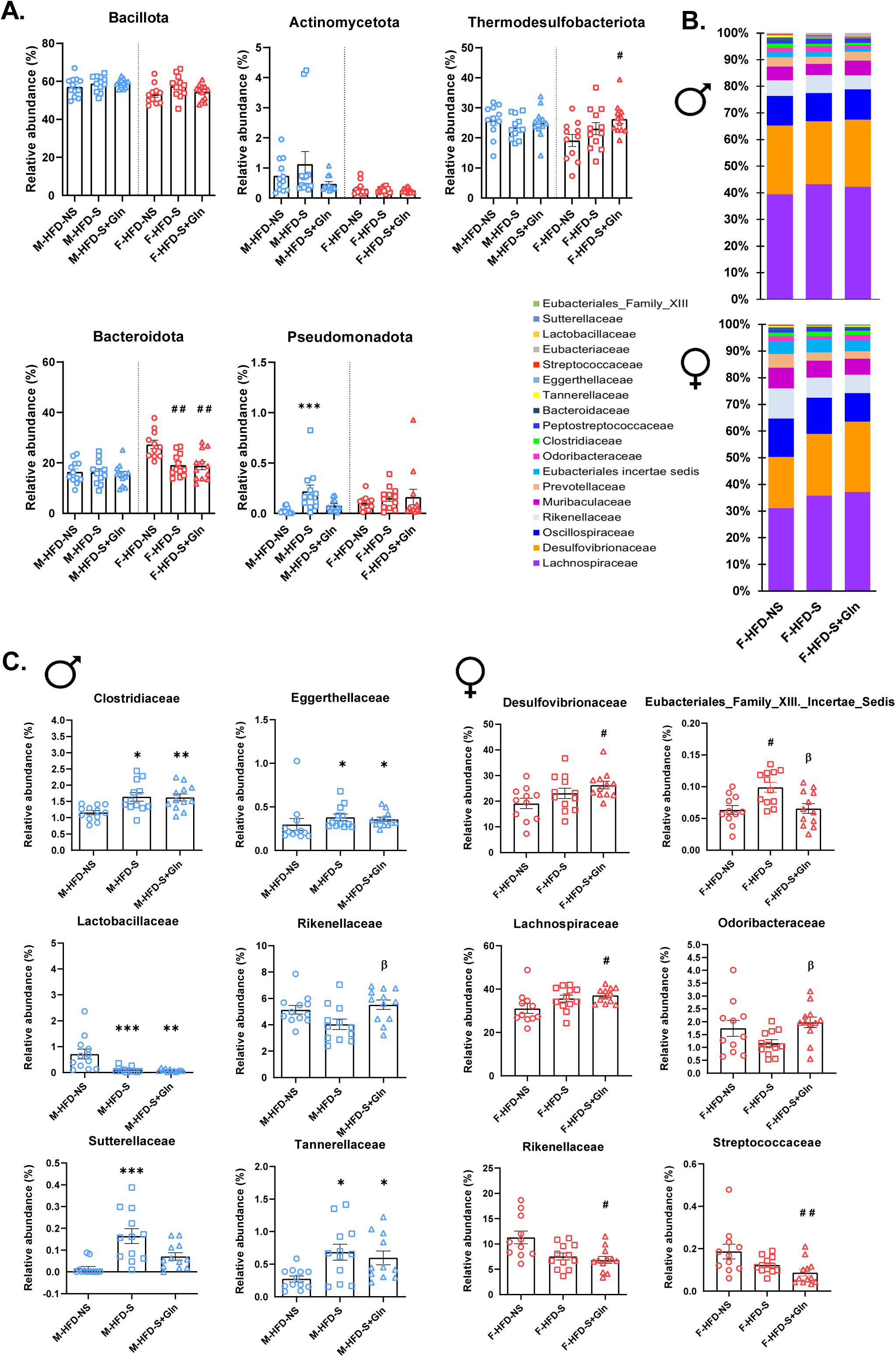
Effects of glutamine (Gln) supplementation on cecal microbiota phylum composition in obese male and female mice subjected to stress. The relative abundance of mainly cecal taxa at the phylum (**A**) and families (**B, C**) levels in male (M) and female (F) mice was assessed at week 14 (W14) fed with high fat diet (HFD) and receiving or not glutamine (Gln) in drinking water from W12 to W14. The last 4 days of the experiments, mice were subjected to a repeated restraint stress (S). Results were compared using the Kruskal-Wallis test followed by Dunn’s post hoc test among groups. Data are expressed as mean ± sem (n=12/group). \**p*<0.05, \*\**p*<0.01, \*\*\**p*<0.001 M-HFD-S or M-HFD-S+Gn *vs* M-HFD-NS; #*p*<0.05, ##*p*<0.01, F-HFD-S or F-HFD-S+Gn *vs* F-HFD-NS; β *p*<0.01 M-ou F-HFD-S *vs* M-ou F-HFD-S+Gln.

**Fig. 6:**
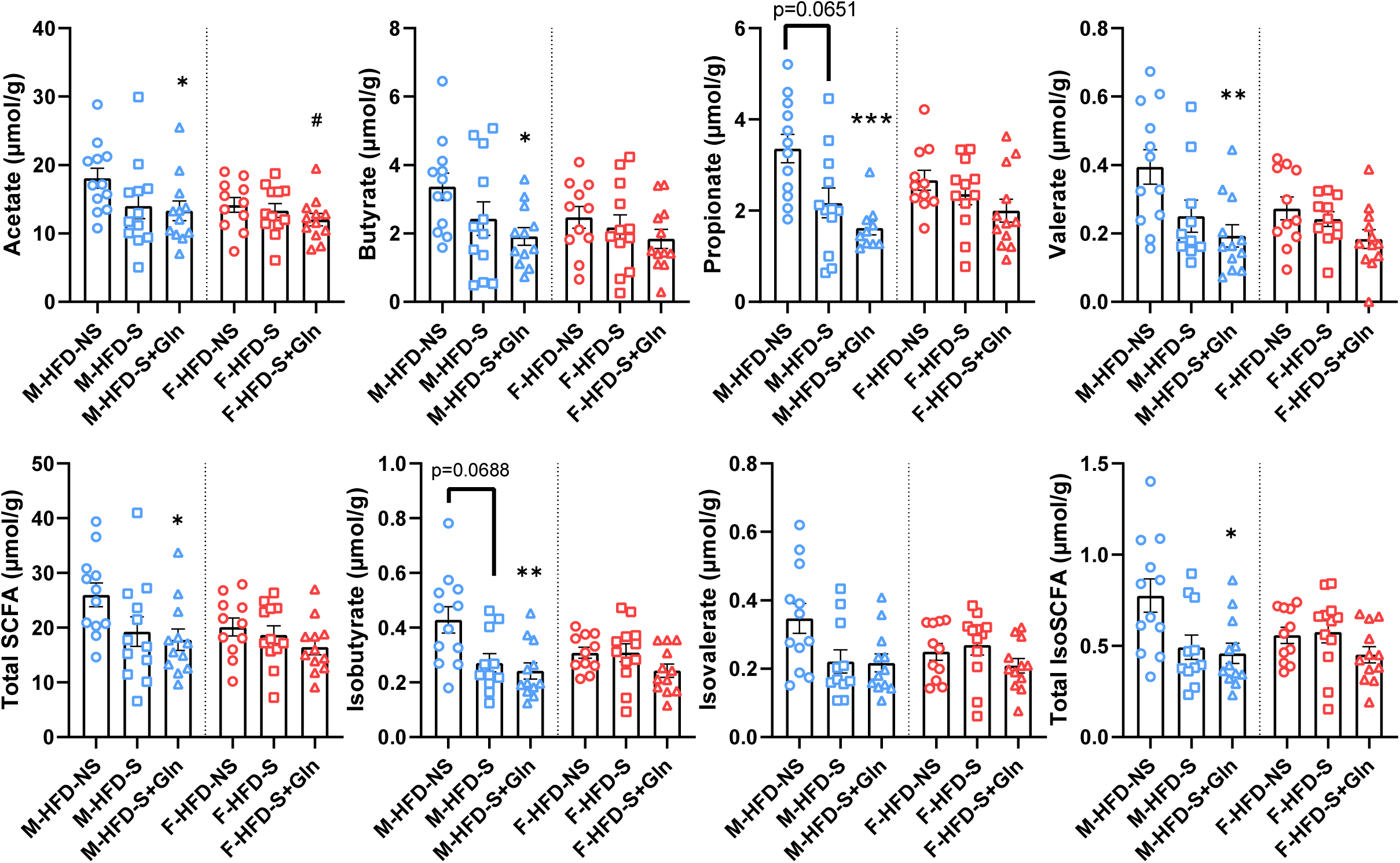
Effects of glutamine (Gln) supplementation on cecal short-chain fatty acids (SCFAs) levels in obese male and female mice subjected to stress. The cecal short-chain fatty acids (**A**) and iso-type short-chain fatty acids (**B**) levels were assessed at week 14 (W14) in male (M) and female (F) mice fed high fat diet (HFD) and receiving or not glutamine (Gln) in drinking water from W12 to W14. The last 4 days of the experiments, mice were subjected to a repeated restraint stress (S). Results were compared by Kruskal-Wallis test followed by Dunnett’s multiple comparison tests. Data are expressed as mean ± sem (n=12/group). \**p*<0.05, \*\**p*<0.01, \*\*\**p*<0.001, M-HFD-S or M-HFD-S+Gn vs M-HFD-NS; #*p*<0.05 F-HFD-S or F-HFD-S+Gn *vs* F-HFD-NS.

According to these data, we observed that Gln supplementation associated to stress mainly changed SCFA in male mice. Indeed, acetate, propionate, butyrate, valerate, isobutyrate and total SCFA were reduced in Gln-supplemented stressed HFD male mice compared to unstressed HFD mice (Fig. 6). In female mice, only acetate level was reduced.

## Discussion

Mood disorders and functional digestive disorders are more frequent in patients with class III obesity compared to the general population (29,30). Psychological stress is an important trigger of these comorbidities, more frequently observed in females. Bridgewater *et al.* highlighted the importance of considering sex as a biological variable in studies on the role of gut microbiota in obesity-related disorders (31). Interestingly, previous data suggest that Gln may be a good candidate to limit stress-induced intestinal disorders (21,23) or obesity-associated metabolic disorders (15,18,32). In the present paper, we report that chronic restraint stress induced sex-dependent effects on gut microbiota, intestinal and metabolic responses in mice with diet-induced obesity. In addition, Gln supplementation also influenced the stress-response in a sex-dependent manner.

In the present work, we used the chronic restraint stress model because of a previous study reporting increase of intestinal permeability (22), a common pathophysiological mechanism described in IBS, anxiety-like disorders and obesity. Even if chronic restraint stress induced an increase of plasma corticosterone in obese mice from both sexes, male and female mice exhibited different stress responses considering body weight and composition, metabolic markers or gut microbiota. Indeed, male mice showed a more pronounced loss of body weight and lean mass than females. In addition, stress ameliorated in males, but not females, the glycemia control by limiting glucose intolerance, by improving fasting glycemia and plasma resistin and by limiting IL-6 mRNA expression in the perigonadal adipose tissue. In accordance with our results, Hatton-Jones *et al.* observed similar effects of chronic restraint stress on glucose metabolism in male mice under western diet with a reduction of fasting glycemia and improvement of glucose tolerance test response (33) but did not enroll female mice in the study. We thus report, for the first time, the sex-specific glycaemic response to stress in obese mice.

Stress was also associated with sex-dependent modifications of cecal microbiota composition, while intestinal permeability remained unchanged in both sexes. By using the water avoidance stress model, we were able to show that stress exacerbated gut barrier disruption in obese male mice compared to lean male mice (12). However, the colonic expression of mRNA encoding tight junction proteins remained unchanged in male mice under western diet and chronic restrain stress (33), as observed in the present work. Concerning cecal microbiota, we did not show differences in α and β diversities but modifications in the phyla, families and species abundances. Accordingly, after alternative restraint stress and forced swimming stress for two weeks, no changes in α and β diversities but complex interactions between diet, stress and sex had been revealed at the phylum or family level (34). In obese males, *Pseudomonadota* abundance was increased in response to stress, while female mice exhibited a decrease of *Bacteroidota* abundance. At the family level, the abundance of *Eggerthellaceae*, *Tannerellaceae* and *Clostridiaceae* was increased in response to stress in male obese mice, while *Lactobacillaceae* showed a lower abundance. In obese females, only *Eubacteriales_Family_XII_Incertae_Sedis* abundance was affected by stress. These data are in accordance with previous data showing an increase of *Pseudomonadota* in stressed male mice (33,35). By combining maternal immune activation, maternal separation and maternal unpredictable chronic mild stress, Rincel *et al.* showed major changes in the gut microbiota composition in male mice compared to limited modifications in female mice (36), as observed in the present study. Similarly, Brix *et al.* recently reported that maternal high fat diet induced stronger effects on gut microbiota in male than in female offspring in response to early life stress (37). It should be thus interesting to decipher the mechanisms involved in this sex-dependent response and to know the contribution of sexual hormones, but also gut microbiota and adipocyte cortisol metabolism, as previously suggested (38).

In the present paper, we also evaluated the effects of Gln supplementation. Pilot studies suggested that Gln supplementation may improve body weight and insulin resistance in patients with overweight or obesity (16–18). In addition, a reduction of Gln content in adipose tissue has been reported during obesity that was associated to low-grade inflammation and insulin-resistance (32). During stress, Gln revealed beneficial properties by limiting intestinal permeability and inflammatory state (21–23) and by limiting anxiety-like and depressive disorders (39). Finally, Gln also showed beneficial effects in patients suffering of irritable bowel syndrome in two randomized controlled trials (23,40). In the present study, Gln partly reduced in stressed female mice, but not in male mice, the plasma corticosterone that was associated with a limitation of body weight loss and an increase of plasma adiponectin. In the previous animal studies (21,22,39), Gln supplementation was not associated with corticosterone changes but only male mice have been studied. By contrast, both male and female mice supplemented with Gln had an improvement of colonic permeability measured in Ussing chambers in accordance with previous data in lean male mice (21,22). To our knowledge, we report for the first time the impact of Gln supplementation on cecal microbiota in obese stressed mice. We recently described the effects of Gln on cecal microbiota in unstressed obese mice (15) showing that Gln changed the cecal microbiota composition in females but not in males. In the present study, Gln prevented the increase of the abundance of *Pseudomonadota* phylum and the *Sutterellaceae* family in stressed obese males. In addition, in males, Gln supplementation was associated with a significant decrease of SCFAs. These data suggest that Gln has stronger effects on gut microbiota in males than in females under stressed conditions that are not associated with synergistic effects on glucose metabolism for instance. Further investigation should decipher the mechanisms involved in the Gln effects and should evaluate the impact of Gln administered in early stages of obesity.

In conclusions, our study reveals sex-dependent alterations in response to stress and Gln supplementation in mice with diet-induced obesity. Stress has beneficial effects on glycemia control in male mice without additive effects of Gln supplementation. By contrast, Gln reduces stress-induced corticosterone level and body weight loss only in females. These data, as well as the differential impact of Gln on intestinal permeability and gut microbiota according to the sex, deserves further investigations to decipher the underlying mechanisms.

## Ethics statement

The protocol was approved by the regional ethics committee (authorization on APAFIS #29283-2021012114574889 v5) and performed in accordance with the current French and European regulations.

## Acknowledgements

We thank Pamela Lecras and Dr David Vaudry from the PRIMACEN platform (HeRaCleS Inserm US51, CNRS UAR 2026, Université de Rouen Normandie, France) and the staff of GenoToul platform (Toulouse, France).

## Funding

The present study was supported by the French Agency for Research (OBEGLU, ANR-20-CE17-0012), the European Society for Clinical Nutrition and Metabolism (ESPEN Fellowship), by the Nutricia Research Foundation, and by European Union and Normandie Regional Council. Europe gets involved in Normandie with European Regional Development Fund (ERDF). CL received the support of the university of Rouen Normandy during her PhD and VD from the Inserm and Normandie Regional council. These funders did not participate in the design, implementation, analysis, and interpretation of the data.

## Author’s contribution

Conceptualization, CL, AG, PD, VDo and MC; CL, AT, MM, AG and MC, formal analysis; CL, AT, CE-B, VDr, CBF, CG, JB, LL, MM, VDo and AG, investigation; CL, VDo, AG and MC, original draft preparation; writing review and editing, AG, VDo and MC.

## Figures legends

**Supplemental Fig. S1:**
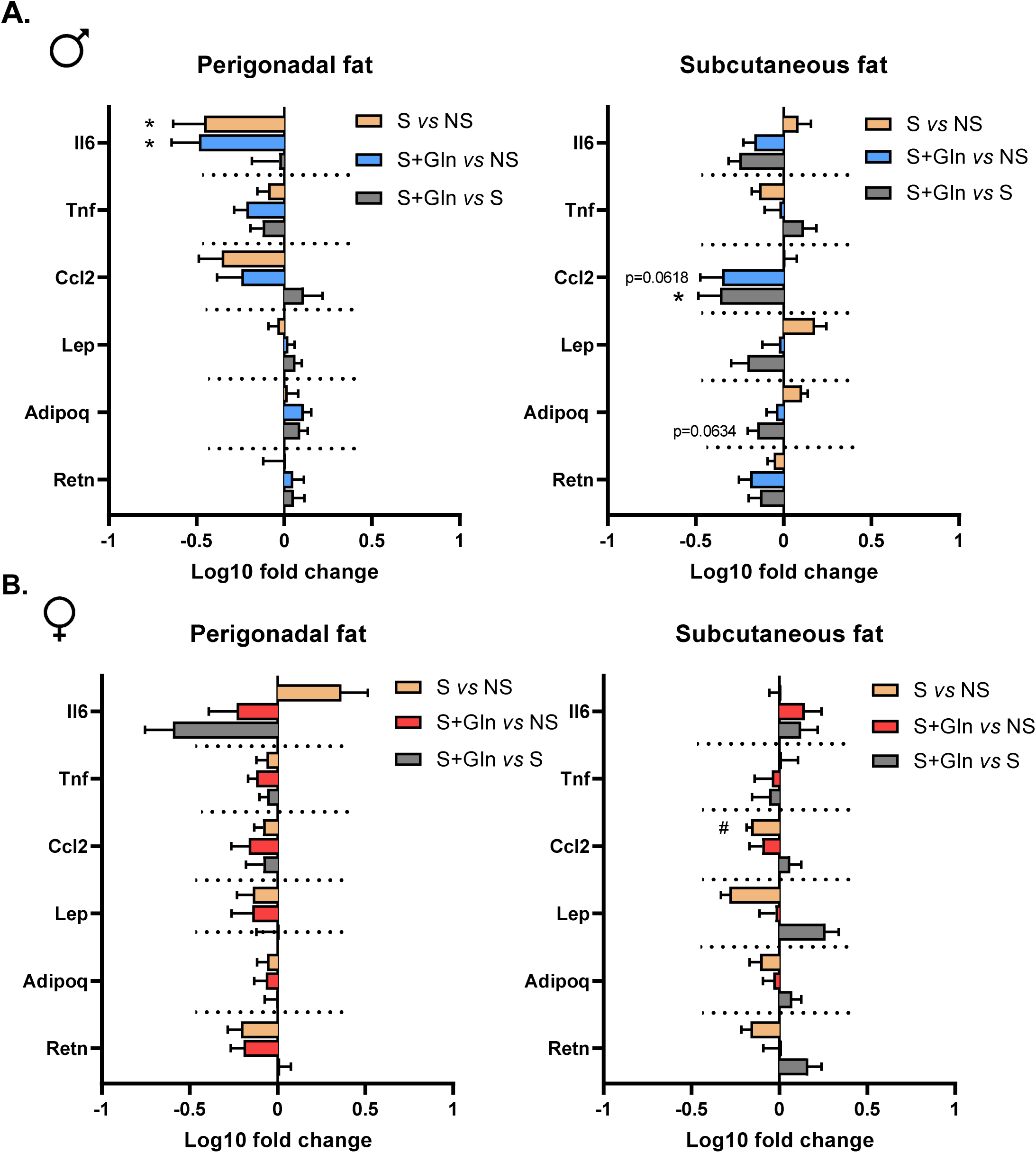
Effects of glutamine (Gln) supplementation on white adipose tissue in obese male and female mice subjected to stress. The levels of mRNA encoding for *Il6* (Interleukin 6), *Tnf* (Tumor necrosis factor alpha), *Ccl2* (C-C motif Chemokine ligand 2 or MCP1), *Lep* (Leptin), *Retn* (Resistin) and *Adipoq* (adiponectin) were measured in perigonadal and subcutaneous fat at week 14 (W14) in male (**A**) and female (**B**) mice fed with high fat diet (HFD) and receiving or not glutamine (Gln) in drinking water from W12 to W14. The last 4 days of the experiments, mice were subjected to a repeated restraint stress (S). Results were compared by Kruskal-Wallis test followed by Dunnett’s multiple comparison tests. Data are presented as ratio log10 [(HFD-S or HFD-S+Gln) / mean HFD-NS] and expressed as mean ± sem (n=12/group). \**p*<0.05 M-HFD-S or M-HFD-S+Gn *vs* M-HFD-NS; #*p*<0.05 F-HFD-S or F-HFD-S+Gn *vs* F-HFD-NS.

**Supplemental Fig. S2:**
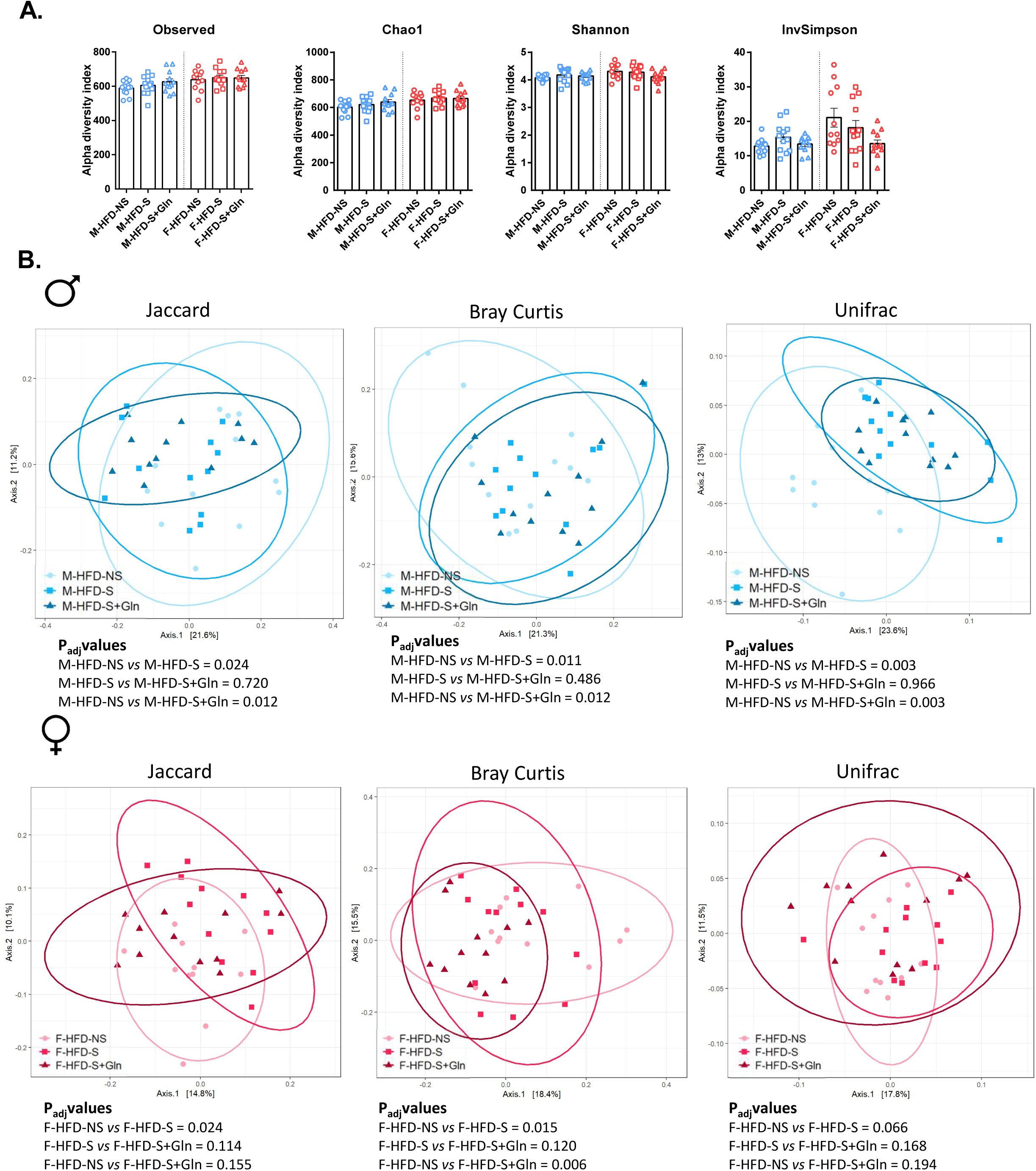
Effects of glutamine (Gln) supplementation on alpha and beta diversity of cecal bacteria species in obese male and female mice subjected to stress. Analysis was based on 16S rDNA sequencing of the cecum content. Observed species richness, Chaos, Shannon and Inverse Simpson Index as indicators of α-diversity (**A**). Principal coordinates analysis (PCoA) of, Jaccard, Bray-Curtis and Unifrac compositional dissimilarity at the amplicon sequence variant (ASV) level (**B**). Each dot represents of one mouse. For α-diversity index values are means ± sem. Results were compared by Kruskal-Wallis test followed by Dunnett’s multiple comparison tests. For all, a value of *p*<0.05 was considered significant.

**Supplemental Fig. S3:**
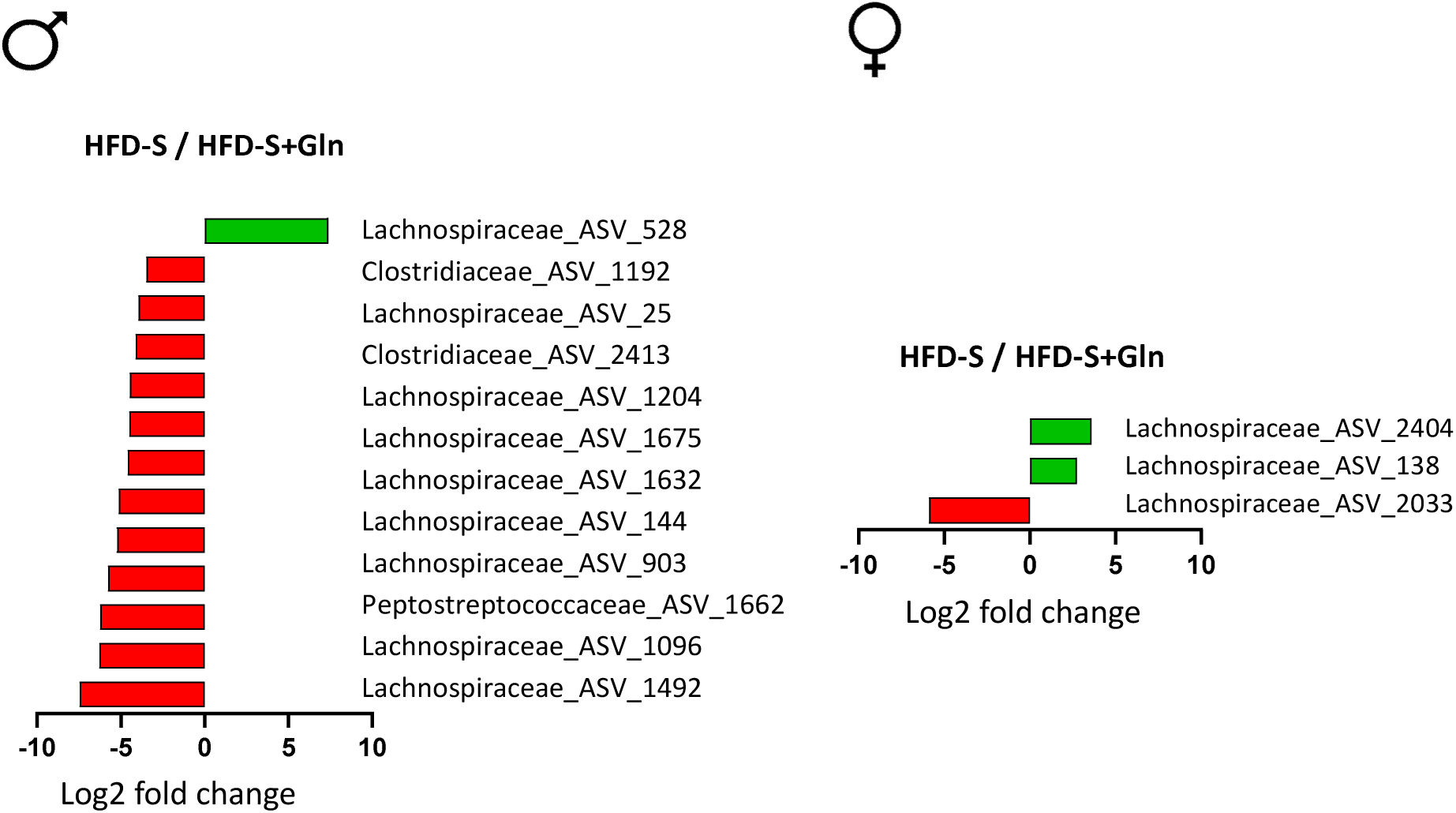
Effects of glutamine (Gln) supplementation on cecal bacteria species in obese male and female mice subjected to stress. Graphic representation of differentially abundant ASVs (with the family to which they belong) between HFD-S and HFD-S-Gln in male and female mice with a logarithmic scale (log-2) used for the x-axis. Data are presented as ratio log2 [mean HFD-S / mean HFD-S+Gln]. Only data with ratio log2 Fold change > 2, a significant (*p*<0.05) effect size and a high relative abundance (more than 30 copies per sample in at least 25% of the samples) are represented.

## Notes

### Competing Interest Statement

The authors have declared no competing interest.

## References

1. Marques A, Peralta M, Naia A, Loureiro N, de Matos MG. Prevalence of adult overweight and obesity in 20 European countries, 2014. Eur J Public Health. 01 2018;28(2):295–300.

2. Ward ZJ, Bleich SN, Cradock AL, Barrett JL, Giles CM, Flax C, et al. Projected U.S. State-Level Prevalence of Adult Obesity and Severe Obesity. N Engl J Med. 19 déc 2019;381(25):2440–50.

3. Aron-Wisnewsky J, Warmbrunn MV, Nieuwdorp M, Clément K. Metabolism and Metabolic Disorders and the Microbiome: The Intestinal Microbiota Associated With Obesity, Lipid Metabolism, and Metabolic Health-Pathophysiology and Therapeutic Strategies. Gastroenterology. janv 2021;160(2):573–99.

4. Cani PD, Bibiloni R, Knauf C, Waget A, Neyrinck AM, Delzenne NM, et al. Changes in gut microbiota control metabolic endotoxemia-induced inflammation in high-fat diet-induced obesity and diabetes in mice. Diabetes. juin 2008;57(6):1470–81.

5. Nascimento JC, Matheus VA, Oliveira RB, Tada SFS, Collares-Buzato CB. High-Fat Diet Induces Disruption of the Tight Junction-Mediated Paracellular Barrier in the Proximal Small Intestine Before the Onset of Type 2 Diabetes and Endotoxemia. Dig Dis Sci. oct 2021;66(10):3359–74.

6. de La Serre CB, Ellis CL, Lee J, Hartman AL, Rutledge JC, Raybould HE. Propensity to high-fat diet-induced obesity in rats is associated with changes in the gut microbiota and gut inflammation. Am J Physiol Gastrointest Liver Physiol. août 2010;299(2):G440–448.

7. Bona MD, Torres CH de M, Lima SCVC, Morais AH de A, Lima AÂM, Maciel BLL. Intestinal Barrier Permeability in Obese Individuals with or without Metabolic Syndrome: A Systematic Review. Nutrients. 3 sept 2022;14(17):3649.

8. Pickett-Blakely O. Obesity and irritable bowel syndrome: a comprehensive review. Gastroenterol Hepatol (N Y). juill 2014;10(7):411–6.

9. Carter D, Beer-Gabel M, Tzur D, Levy G, Derazne E, Novis B, et al. Predictive factors for the diagnosis of irritable bowel syndrome in a large cohort of 440,822 young adults. J Clin Gastroenterol. avr 2015;49(4):300–5.

10. Melchior C, Hreinsson JP, Tack J, Keller J, Aziz Q, Palsson OS, et al. Disorders of the gut-brain interaction among European people with obesity: Prevalence and burden of compatible symptoms. United European Gastroenterol J. juill 2025;13(6):917–28.

11. Bertiaux-Vandaële N, Youmba SB, Belmonte L, Lecleire S, Antonietti M, Gourcerol G, et al. The expression and the cellular distribution of the tight junction proteins are altered in irritable bowel syndrome patients with differences according to the disease subtype. Am J Gastroenterol. déc 2011;106(12):2165–73.

12. Bahlouli W, Breton J, Lelouard M, L’Huillier C, Tirelle P, Salameh E, et al. Stress-induced intestinal barrier dysfunction is exacerbated during diet-induced obesity. J Nutr Biochem. 2020;81:108382.

13. Pugliese G, Muscogiuri G, Barrea L, Laudisio D, Savastano S, Colao A. Irritable bowel syndrome: a new therapeutic target when treating obesity? Hormones (Athens). déc 2019;18(4):395–9.

14. Wang B, Wu G, Zhou Z, Dai Z, Sun Y, Ji Y, et al. Glutamine and intestinal barrier function. Amino Acids. oct 2015;47(10):2143–54.

15. Lefebvre Candice, Tiffay Adam, Breemeersch Charles-Edward, Dreux Virginie, Bôle-Feysot Christine, Guérin Charlène, et al. Sex-dependent effects of a glutamine supplementation on metabolic disorders, intestinal barrier function and gut microbiota in mice with diet-induced obesity.

16. Opara EC, Petro A, Tevrizian A, Feinglos MN, Surwit RS. L-glutamine supplementation of a high fat diet reduces body weight and attenuates hyperglycemia and hyperinsulinemia in C57BL/6J mice. J Nutr. janv 1996;126(1):273–9.

17. Laviano A, Molfino A, Lacaria MT, Canelli A, De Leo S, Preziosa I, et al. Glutamine supplementation favors weight loss in nondieting obese female patients. A pilot study. Eur J Clin Nutr. nov 2014;68(11):1264–6.

18. Abboud KY, Reis SK, Martelli ME, Zordão OP, Tannihão F, de Souza AZZ, et al. Oral Glutamine Supplementation Reduces Obesity, Pro-Inflammatory Markers, and Improves Insulin Sensitivity in DIO Wistar Rats and Reduces Waist Circumference in Overweight and Obese Humans. Nutrients. 1 mars 2019;11(3).

19. de Souza AZZ, Zambom AZ, Abboud KY, Reis SK, Tannihão F, Guadagnini D, et al. Oral supplementation with L-glutamine alters gut microbiota of obese and overweight adults: A pilot study. Nutrition. juin 2015;31(6):884–9.

20. Moloney RD, O’Mahony SM, Dinan TG, Cryan JF. Stress-induced visceral pain: toward animal models of irritable-bowel syndrome and associated comorbidities. Front Psychiatry. 2015;6:15.

21. Ghouzali I, Lemaitre C, Bahlouli W, Azhar S, Bôle-Feysot C, Meleine M, et al. Targeting immunoproteasome and glutamine supplementation prevent intestinal hyperpermeability. Biochim Biophys Acta Gen Subj. janv 2017;1861(1 Pt A):3278–88.

22. Langlois LD, Oddoux S, Aublé K, Violette P, Déchelotte P, Noël A, et al. Effects of Glutamine, Curcumin and Fish Bioactive Peptides Alone or in Combination on Intestinal Permeability in a Chronic-Restraint Stress Model. Int J Mol Sci. 13 avr 2023;24(8):7220.

23. Zhou Q, Verne ML, Fields JZ, Lefante JJ, Basra S, Salameh H, et al. Randomised placebo-controlled trial of dietary glutamine supplements for postinfectious irritable bowel syndrome. Gut. 2019;68(6):996–1002.

24. Medrikova D, Jilkova ZM, Bardova K, Janovska P, Rossmeisl M, Kopecky J. Sex differences during the course of diet-induced obesity in mice: adipose tissue expandability and glycemic control. Int J Obes (Lond). févr 2012;36(2):262–72.

25. Elzinga SE, Savelieff MG, O’Brien PD, Mendelson FE, Hayes JM, Feldman EL. Sex differences in insulin resistance, but not peripheral neuropathy, in a diet-induced prediabetes mouse model. Dis Model Mech. 1 avr 2021;14(4):dmm048909.

26. Lefebvre C, Tiffay A, Breemeersch CE, Dreux V, Bôle-Feysot C, Guérin C, et al. Sex-dependent effects of a high fat diet on metabolic disorders, intestinal barrier function and gut microbiota in mouse. Sci Rep. 27 août 2024;14(1):19835.

27. McMurdie PJ, Holmes S. phyloseq: An R Package for Reproducible Interactive Analysis and Graphics of Microbiome Census Data. PLOS ONE. 22 avr 2013;8(4):e61217.

28. Love MI, Huber W, Anders S. Moderated estimation of fold change and dispersion for RNA-seq data with DESeq2. Genome Biol. 2014;15(12):550.

29. Schneck AS, Anty R, Tran A, Hastier A, Amor IB, Gugenheim J, et al. Increased Prevalence of Irritable Bowel Syndrome in a Cohort of French Morbidly Obese Patients Candidate for Bariatric Surgery. Obes Surg. 2016;26(7):1525–30.

30. Gallagher C, Waidyatillake N, Pirkis J, Lambert K, Cassim R, Dharmage S, et al. The long-term effects of childhood adiposity on depression and anxiety in adulthood: A systematic review. Obesity (Silver Spring). sept 2023;31(9):2218–28.

31. Bridgewater LC, Zhang C, Wu Y, Hu W, Zhang Q, Wang J, et al. Gender-based differences in host behavior and gut microbiota composition in response to high fat diet and stress in a mouse model. Sci Rep. 7 sept 2017;7(1):10776.

32. Petrus P, Lecoutre S, Dollet L, Wiel C, Sulen A, Gao H, et al. Glutamine Links Obesity to Inflammation in Human White Adipose Tissue. Cell Metab. 4 févr 2020;31(2):375–390.e11.

33. Hatton-Jones KM, du Toit EF, Cox AJ. Effect of chronic restraint stress and western-diet feeding on colonic regulatory gene expression in mice. Neurogastroenterol Motil. avr 2022;34(4):e14300.

34. Lyte JM, Koester LR, Daniels KM, Lyte M. Distinct Cecal and Fecal Microbiome Responses to Stress Are Accompanied by Sex– and Diet-Dependent Changes in Behavior and Gut Serotonin. Front Neurosci. 2022;16:827343.

35. Wang M, Sun P, Li Z, Li J, Lv X, Chen S, et al. Eucommiae cortex polysaccharides attenuate gut microbiota dysbiosis and neuroinflammation in mice exposed to chronic unpredictable mild stress: Beneficial in ameliorating depressive-like behaviors. J Affect Disord. 1 août 2023;334:278–92.

36. Rincel M, Aubert P, Chevalier J, Grohard PA, Basso L, Monchaux de Oliveira C, et al. Multi-hit early life adversity affects gut microbiota, brain and behavior in a sex-dependent manner. Brain Behav Immun. août 2019;80:179–92.

37. Brix LM, Monleon D, Collado MC, Ederveen THA, Toksöz I, Bordes J, et al. Metabolic effects of early life stress and pre-pregnancy obesity are long lasting and sex specific in mice. Eur J Neurosci. juill 2023;58(1):2215–31.

38. Incollingo Rodriguez AC, Epel ES, White ML, Standen EC, Seckl JR, Tomiyama AJ. Hypothalamic-pituitary-adrenal axis dysregulation and cortisol activity in obesity: A systematic review. Psychoneuroendocrinology. déc 2015;62:301–18.

39. Faucher P, Dries A, Mousset PY, Leboyer M, Dore J, Beracochea D. Synergistic effects of Lacticaseibacillus rhamnosus GG, glutamine, and curcumin on chronic unpredictable mild stress-induced depression in a mouse model. Benef Microbes. 3 août 2022;13(3):253–64.

40. Rastgoo S, Ebrahimi-Daryani N, Agah S, Karimi S, Taher M, Rashidkhani B, et al. Glutamine Supplementation Enhances the Effects of a Low FODMAP Diet in Irritable Bowel Syndrome Management. Front Nutr. 2021;8:746703.

